# EpiDope: A Deep neural network for linear B-cell epitope prediction

**DOI:** 10.1101/2020.05.12.090019

**Authors:** Maximilian Collatz, Florian Mock, Martin Hölzer, Emanuel Barth, Konrad Sachse, Manja Marz

## Abstract

By binding to specific structures on antigenic proteins, the so-called epitopes, B-cell antibodies can neutralize pathogens. The identification of B-cell epitopes is of great value for the development of specific serodiagnostic assays and the optimization of medical therapy. However, identifying diagnostically or therapeutically relevant epitopes is a challenging task that usually involves extensive laboratory work. In this study, we show that the time, cost and labor-intensive process of epitope detection in the lab can be significantly shortened by using *in silico* prediction. Here we present `EpiDope`, a python tool which uses a deep neural network to detect B-cell epitope regions on individual protein sequences (github.com/mcollatz/EpiDope). With an area under the curve (AUC) between 0.67 ± 0.07 in the ROC curve, `EpiDope` exceeds all other currently used B-cell prediction tools. Moreover, for AUC10% (AUC for a false-positive rate < 0.1), `EpiDope` improves the prediction accuracy in comparison to other state-of-the-art methods. Our software is shown to reliably predict linear B-cell epitopes of a given protein sequence, thus contributing to a significant reduction of laboratory experiments and costs required for the conventional approach.

## Introduction

The public health system is highly dependent on the use of vaccines to protect the population from a range of dangerous infectious diseases. Through decades of systematic vaccination, diseases like measles, mumps, rubella, pertussis, poliomyelitis, diphtheria, tetanus and others have been largely eradicated [1, 2]. Vaccination is also an efficient approach to avoid or reduce prescriptions of antibiotics and, as a consequence, minimize the emergence of ever more multi-resistant strains of microbial pathogens. To assess the degree of protection of vaccination at population level, faster and more efficient serological tools need to be developed. They should be capable of identifying geographical and social heterogeneities in the diversity of population immunity [3, 4]. In addition, it is important to know the status of a patient’s immunization in order to avoid unnecessary vaccinations. In cases where this is not or only incompletely documented, various tests can be used to determine which specific immunities already exist and which vaccinations are missing. The serological tests currently used, however, are still largely based on ELISA technology and are only able to detect antibodies against a single, specific pathogen [4]. These tests are not only slow and expensive, but usually also use whole cell antigens to detect antibodies, which limits their specificity [5].

B-cell antibodies of the immune system of a host are able to detect certain exposed amino acids and subsequently bind the corresponding antigenic proteins. These bound protein regions are called epitopes and represent the interface between infection and immune response [6, 7]. The antibody part that binds the epitope is called paratope. Epitopes themselves are not intrinsic features of a protein, but rather relational units defined by the interaction with a binding paratope [6]. This relatively vague definition makes it a challenging task to predict epitopes *in silico* [7, 8]. Furthermore, epitopes are divided into linear and conformational epitopes, where linear epitopes consist of a contiguous piece of amino acids and conformational epitopes consist of atoms of surface residues that come together by protein folding [6]. In this study, we will focus on the prediction of linear B-cell epitopes.

A frequently used tool to predict linear B-cell epitopes is `BepiPred2` [9]. Jespersen *et al.* state an area under the curve (AUC) of 0.57 for their tool’s receiver operating characteristic (ROC) curve. This is on par with other prediction scales provided by the “Immune Epitope Database” IEDB (http://tools.iedb.org/main/bcell/) and demonstrates the difficulty of *in silico* epitope identification. Therefore, we developed `EpiDope`, a tool based on deep neural networks (DNN) to detect epitopic regions in proteins based on their primary amino acid sequence.

DNNs are often used in complex classification problems with limited knowledge about useful features of the objects to be classified. With sufficient data, a DNN can automatically recognize appropriate classification features, making DNNs very suitable for the prediction of linear B-Cell epitopes [10].

We will show that our DNN-based program `EpiDope` succeeds in identifying linear B-cell epitopes with a ROC AUC of 0.67 ± 0.07, which significantly exceeds previous methods. This helps to considerably reduce the number of potential linear epitopes to be validated experimentally and, thus, can accelerate the development of serological assays and immunotherapeutic approaches.

## Material and Methods

### Data

The “IEDB Linear Epitope Dataset” (available at http://www.cbs.dtu.dk/services/BepiPred/download.php), which was also used for the evaluation of the B-cell epitope prediction tool `BepiPred2` [9], served as a training basis for our DNN. It consists of 30,556 protein sequences, in which each sequence contains a marked region, in the following called ‘verified regions’, that represents an experimentally verified epitope or non-epitope. The subset of epitopes has a average length of 13.99, whereas the subset of non-epitopes has a average length of 13.20 (see Table 1).

**Table 1:** Comparison of the original dataset provided by Jespersen et al.[9], and the redundancy reduced data used as training data.

|  | original | reduced |
| --- | --- | --- |
| # verified regions | 30,556 | 24,610 |
| # epitopes | 11,834 | 8,519 |
| # non-epitopes | 18,722 | 16,091 |
| median length | 15 | 15 |
| avg. length | 13.50 | 13.27 |
| avg. length epitope | 13.99 | 13.88 |
| avg. length non-epitope | 13.20 | 12.95 |

### Data preparation

In order to ensure the best possible training basis for the DNN, we pre-processed the dataset in several steps (see Figure 1). First, we merged identical protein sequences while keeping the information about their verified regions, resulting in a reduced dataset containing 3,158 proteins preserving all 30,556 verified regions (Figure 1 A and B). Second, to reduce redundancy by non-identical but highly similar protein sequences, we clustered all sequences with `cd-hit` [11] using an identity threshold of 0.8 (Figure 1 C). This resulted in 1,798 protein sequence clusters. From each cluster, only the protein sequence containing the largest number of verified regions was retained, reducing the number of verified regions by 19.46 % to 24610 (see Table 1, Figure 1 D). This reduces the number of very similar and thus overrepresented proteins in the data, as these might bias the training of the DNN.

**Figure 1:**
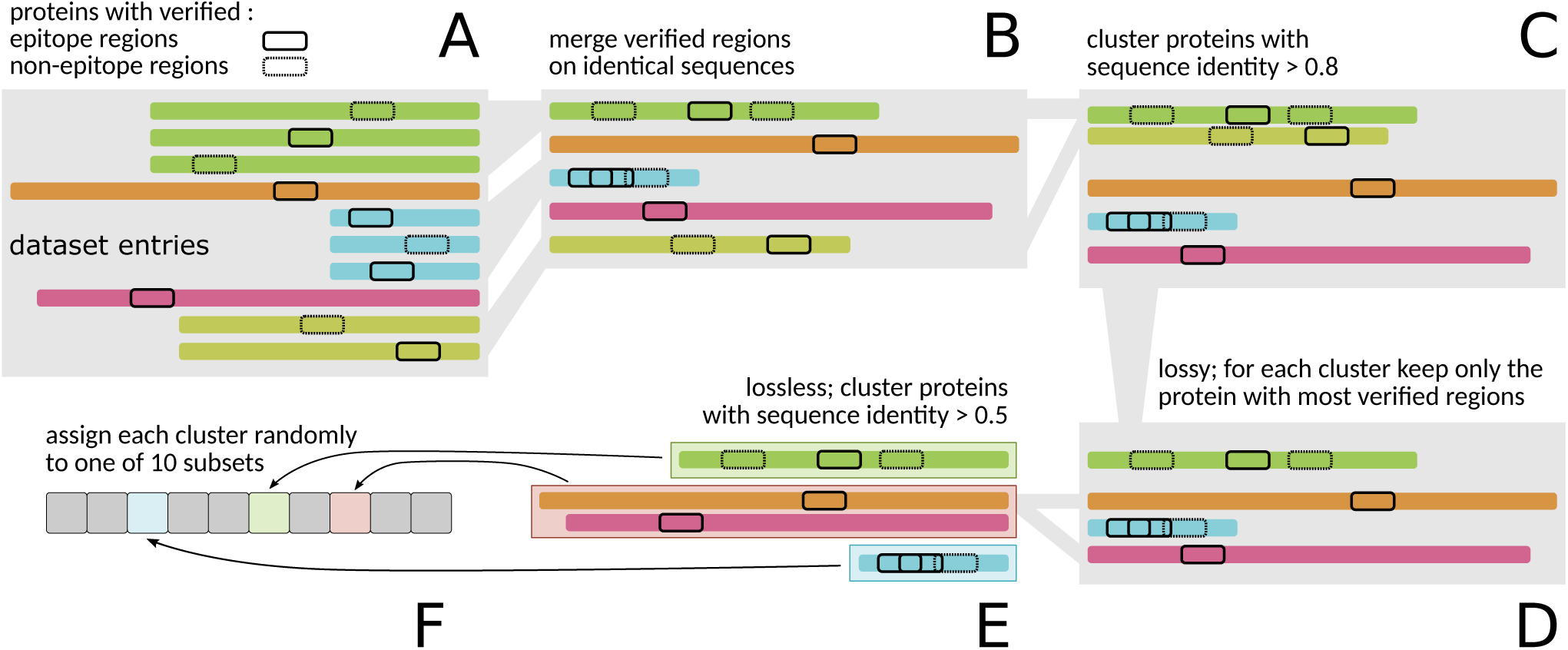
The data preparation to generate the training and validation set. First all identical amino acid sequences from the raw dataset (A) are merged while preserving the verified regions (B). In the next step (C), all remaining sequences are clustered with a sequence identity of 0.8 and higher. For each of the resulting clusters only the sequence with the highest number of verified regions is selected (D). These selected sequences are then clustered again, this time with a sequence identity of 0.5 and higher (E). Each cluster is then assigned to only one of ten different subsets for the 10-fold cross-validation (F). The data preparation generates training and test data with a low sequence identity of < 0.5 while limiting the loss of potential sequences.

The clustering step was repeated on the reduced protein sequences using an identity threshold of 0.5, resulting in 1,378 sequence clusters. These clusters were then used to build the 10-fold cross-validation for the DNN, where the data was divided into ten equally sized subsets according to the number of clusters. This sequence reduction approach was implemented with the following advantage in mind: The DNN training, and test sets share no proteins with a sequence identity of more than 0.5, ensuring that both sets are as independent as possible from each other, avoiding the DNN from simply memorizing specific epitopes.

### Deep neural network architecture

We compared several DNN architectures, including different ordering, layer types and numbers of nodes. The DNN architecture used in `EpiDope` consists of two parts (see Figure 2). The first part (Figure 2 A) uses context-sensitive embeddings of amino acids produced by an ELMo DNN. This DNN was previously trained by Heinzinger *et al.* to encode various chemical, physical and structural information and was demonstrated to be usable for various high-quality predictions on protein sequences [12]. Each amino acid of a given protein sequence is encoded in a vector of length 1,024 which encodes the chemical, physical and structural information. These embeddings are the input for a bidirectional LSTM layer with 2×5 nodes [13], followed by a dense layer containing ten nodes.

**Figure 2:**
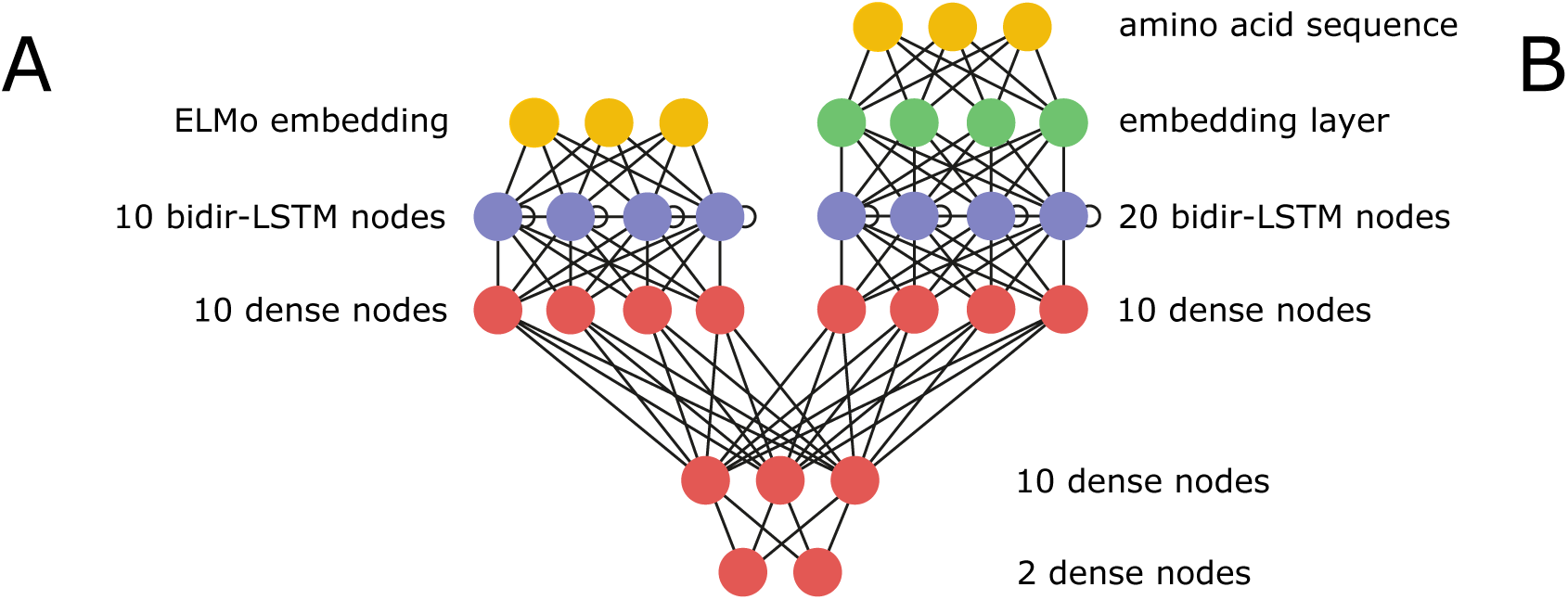
The DNN architecture of `EpiDope` consists of two parts. The first (A) uses context-sensitive ELMo embeddings for the epitope prediction. These embeddings are previously calculated by an ELMo DNN. The second part (B) uses classic embeddings for prediction. These classic embeddings are not context-sensitive. Both parts are then joined to predict two classes, epitopes and non-epitopes.

The second part of our DNN architecture (Figure 2B) encodes each amino acid into a vector of length ten. This embedding is not context-sensitive and is trained together with the rest of the DNN. This embedding layer is connected to a bidirectional LSTM layer with 2×10 nodes, again followed by a dense layer with ten nodes.

Both dense layers are then connected with an additional dense layer containing ten nodes, which is concluded by the output layer with two nodes representing the two classes, *epitope* and *non-epitope*. Note, the number of nodes in this DNN is comparatively low. However, due to the high dimensionality of the context-sensitive embedding (1024 per amino acid), the number of parameters tuned by the DNN is substantial.

### Evaluation of epitope prediction approaches

We compared our tool `EpiDope` against five frequently tools for linear B-cell epitope prediction from the IEDB (http://tools.iedb.org/bcell/) in their latest version. Namely `Bepipred 2.0` [9], `Parker Hydrophilicity prediction` [14], `Chou and Fasman beta turn prediction` [15], `Emini surface accessibility scale` [16], `Kolaskar and Tongaonkar antigenicity scale` [17] and the independent prediction tool for intrinsically unstructured protein regions `IUPred` [18].

For each tool, we calculated the corresponding prediction values for the entire protein sequence on each amino acid and sliced out the verified regions. For each of the sliced regions, we calculated the average score as the prediction score to discriminate between epitope and non-epitope. For the `antigenicity scale` approach we additionally subtracted the mean of the full protein as suggested in the original paper [17].

We compared all the tools using ROC and Precision-Recall curves. For the EpiDope ROC curve, any subset of the 10-fold cross-validation was predicted by the model that did not include this subset in its training data. We calculated one ROC curve per subset. This resulted in ten ROC curves, on which the mean ROC curve was calculated. In the mean ROC curve each of the ten models had an equal influence. The ROC curves of competing tools are calculated on the same data without having to combine multiple predictions, as these tools and scores did not use this data for training.

For the Precision-Recall curve of EpiDope, as with the ROC curve, each subset was predicted by the model that did not use it in the training set. Next, we calculate the Precision-Recall curve on all ten subsets at once rather than calculating ten Precision-Recall curves. Calculating the mean curve from ten Precision-Recall curves could change the balance between the two classes and as such bias the Precision-Recall curve. All other Precision Recall curves were calculated equally.

The ROC curves as well as the Precision-Recall curves were calculated using `scikit-learn` [19].

## Results & Discussion

### The IEDB dataset represents large pathogen variety

In order to achieve a bias-free prediction of epitopes, it is essential to have a large variety of known epitopes from evolutionarily distinct organisms in the training set. Therefore, we initially analyzed the taxonomic origin of the protein sequences provided by the IEDB and visualized the results with the online tool `Pavian metagenomics data explorer` [20], see Figure 3. Since the number of verified regions per protein varies, we analyzed them separately. Our filtered dataset contains 1,798 proteins with 24,610 epitopes and non-epitopes assigned to them.

**Figure 3:**
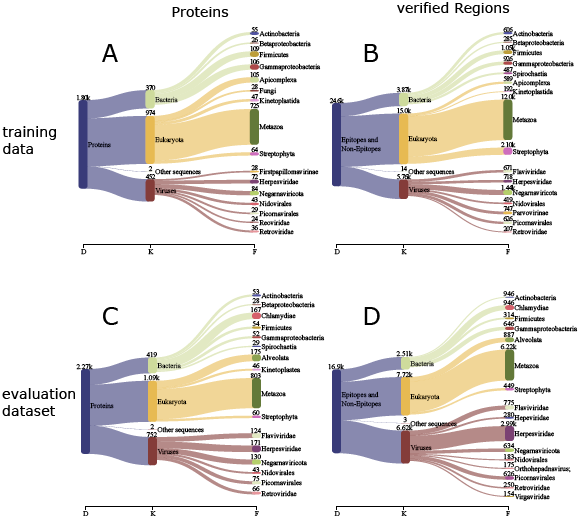
Taxonomic origin of the training data (A and B) and evaluation dataset (C and D). A shows the origin of all 1,798 proteins in the training data, covering a wide variety of kingdoms and families. B shows the taxonomic origin of the epitopes and non-epitopes, with 15.7 % from Bacteria, 61 % Eukaryota and 23.4 % from Viruses. Taxonomic origin of the sequences used in the evaluation dataset: C shows the origin of all 2,267 proteins and D shows the taxonomic origin of the epitopes and non-epitopes, with 14.9 % from Bacteria, 45.7 % from Eukaryota and 39.2 % from Viruses.

At the protein level, the data consist of 16 different families. The four families of Bacteria account for 20.6 %, the five families of Eukaryota for 54.2 %, and the seven families of Viruses for 25.1 % of all proteins in the training data. At the level of the verified regions, we observed a slight shift, so that Eukaryota with 61.0 % are even more pronounced than the Bacteria with 15.7 % and the Viruses with 23.4 %. Overall, the training data display a sufficient degree of taxonomic diversity.

### Evaluation of EpiDope via 10-fold cross-validation

As described in the methods section, we evaluated the performance of `EpiDope` with competing methods on the training data, using 10-fold crossvalidation. Each of the ten subsets consists of proteins that have a sequence identity below 0.5 to all proteins in the other nine subsets (for details, see the ‘Data preparation’ section).

The Receiver operating characteristic (ROC)-curve (Figure 4A) shows that `EpiDope` is with an AUC of 0.67 ± 0.07 clearly outperforming competing prediction approaches, all of which achieved an AUC ≤ 0.56. Despite multiple requests from us to the developers of `BepiPred2`, we could not confirm their stated AUC performance of 0.57 on our reduced and less redundant dataset (see Table 1).

**Figure 4:**
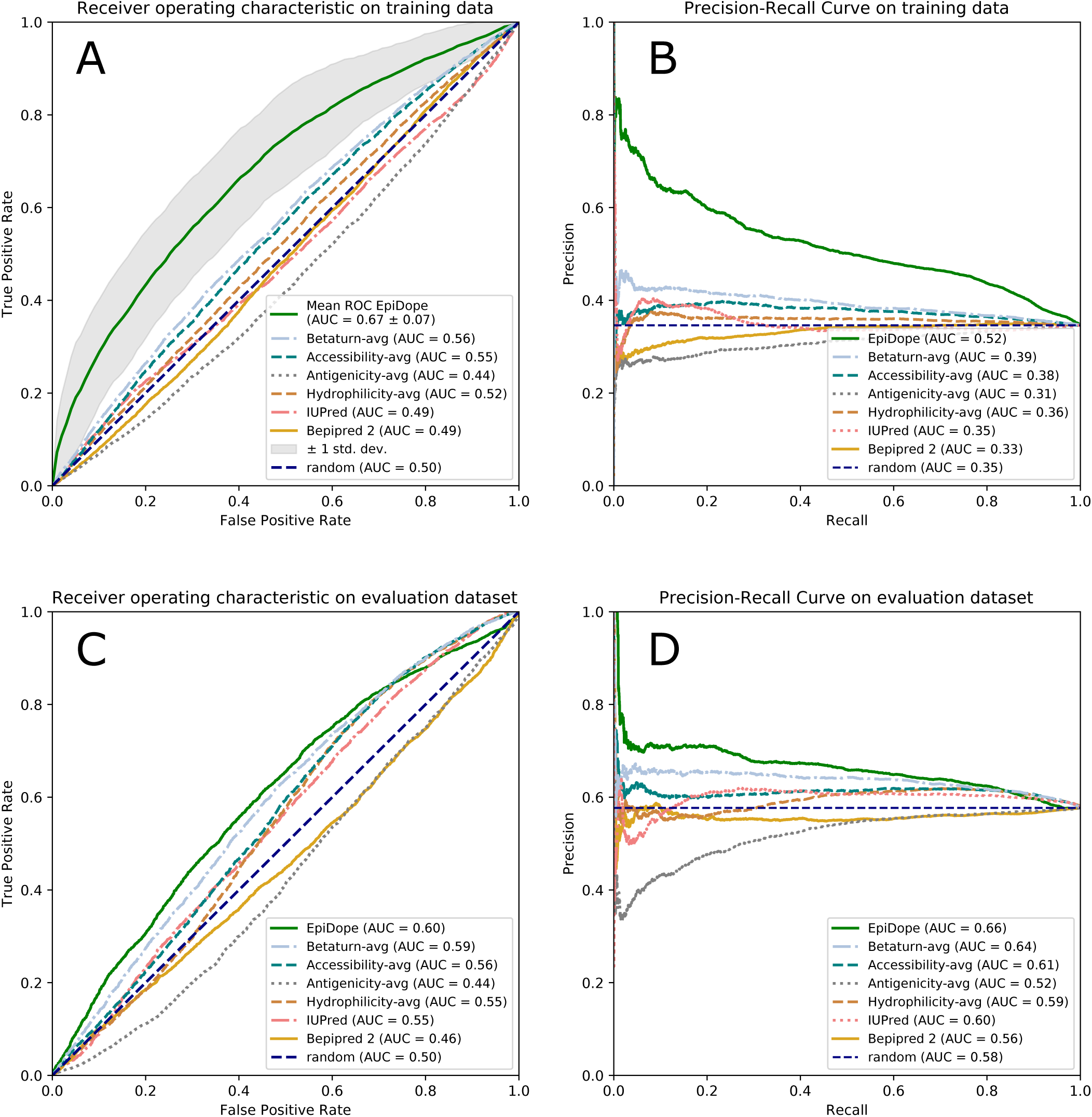
A shows the comparison of the ROC curves between the evaluated tools based on the training dataset. For the mean `EpiDope` ROC curve, every subset of the 10-fold cross-validation was predicted by the model that did not include this subset in its training data. This resulted in ten ROC curves for which the mean ROC curve was calculated (displayed in green) and the standard deviation area (grey). The other ROC curves were calculated on the same data without having to combine multiple predictions, as these tools and scores did not use this data for training. In B, the precision-recall curve shows the trade-off between the number of false positive predictions compared to the number of false negative predictions. This is important as the number of epitopes and non-epitopes are not balanced in the dataset. C shows the ROC curves for the evaluation dataset, which consists of all currently (as of 27.11.2019) well verified regions that are not present in the training data. D shows the precision-recall curve on the evaluation dataset.

Usually, it is not necessary to find all immunodominant epitopes in a proteome. Instead, only a small number of functioning epitopes are required, ideally without having to scan a large number of regions. Therefore the prediction performance in the highest-rated regions is of particular interest. To evaluate the performance of these regions, we used AUC10%. The AUC10% is the AUC of the ROC curve for a False Positive Rate (FPR) < 0.1, normalized, as suggested by the `BepiPred2` developers by multiplying with 10 [9].

`EpiDope` reaches a high AUC10% of 0.151, compared to the second best method (Betaturn) with an AUC10% of 0.070 (see Table 2). Note that Jespersen *et al.* state an AUC10% of 0.08 for `BepiPred2` [9] on the original unreduced dataset. The performance of `EpiDope` relies on the high precision of the top predictions, notable also in the Precision-Recall curve, see Figure 4B.

**Table 2:** Comparison of the AUC and AUC10% of the ROC curve calculated for multiple tools, on the training data. AUC shows the value on the complete ROC curve while AUC10% is the area for a False Positive Rate (FPR) < 0.1 corrected by multiplying with 10. `EpiDope` is compared using 10-fold cross validation.

| training data |  |  |
| --- | --- | --- |
| Tool | AUC | AUC10% |
| EpiDope | 0.670 | 0.151 |
| Betaturn | 0.562 | 0.070 |
| Accessibility | 0.548 | 0.058 |
| Antigenicity | 0.443 | 0.034 |
| Hydrophilicity | 0.521 | 0.053 |
| IUPred | 0.489 | 0.060 |
| Bepipred2 | 0.490 | 0.038 |
| random | 0.500 | 0.050 |

### Evaluation on new data

We created a new dataset that contains all verified regions that were not included in the `BepiPred2` dataset (as of 27.11.2019). All areas that were tested positive by at least two assays were stored as epitopes, whereas all areas tested in at least two assays and not tested positive in any assay were stored as non-epitopes. These are the same conditions that were used to create the `BepiPred2` dataset [9]. Thus this dataset is entirely independent of our training dataset. This second dataset is from now on called evaluation dataset. The evaluation dataset was not pre-processed like the previous dataset (see Figure 1), to preserve a high amount of samples.

In comparison with the training data the evaluation dataset is of comparable size (see Figure 3B and 3D), with the training data having over 24,600 verified regions and the evaluation dataset having over 16,900 verified regions. The evaluation dataset has a lower proportion of eukaryotic verified regions (45.7%) and a higher proportion of viral verified regions (33.1%) compared the training data (eukaryotic 61.0%, viral 23.4%). From the 16 most common families (Figure 3D), five were not present in the training data. Theses families are *Chlamydiae, Alveolata, Hepeviridae, Orthohepadnavirus*, and *Virgaviridae*, representing 5.6 %, %, 1.66 %, 1 % and 0.91 % of the evaluation dataset. The family of *Herpesviridae* is in the evaluation dataset over six times more common, with 17.75 % as compared to the training dataset (2.92 %).

Furthermore, the ratio of epitope regions vs. non-epitope regions changed dramatically, from 35 % epitopes in the training dataset to 58 % in the evaluation dataset.

As with the training dataset, we calculated the predictions of several tools (see ‘Evaluation of epitope prediction approaches’ section for further details) for the verified regions. We calculated the ROC curve and the Precision-Recall curve (see Figure 4C/D) and evaluated the AUC and AUC10% of `EpiDope` and the competing tools (see Table 3).

**Table 3:** Comparison of the AUC and AUC10% of the ROC curve calculated for multiple tools on the evaluation dataset. AUC shows the value on the complete ROC curve, while AUC10% is the area for a false-positive rate (FPR) < 0.1 corrected by multiplying with 10.

| evaluation dataset |  |  |
| --- | --- | --- |
| Tool | AUC | AUC10% |
| EpiDope | 0.605 | 0.089 |
| Betaturn | 0.589 | 0.069 |
| Accessibility | 0.563 | 0.058 |
| Antigenicity | 0.436 | 0.021 |
| Hydrophilicity | 0.550 | 0.048 |
| IUPred | 0.550 | 0.040 |
| Bepipred2 | 0.465 | 0.048 |
| random | 0.500 | 0.050 |

The ROC-curve (see Figure 4C) shows that `EpiDope` is, with an AUC of 0.60, surpassing competing prediction approaches. We can observe a distinctly different result for nearly all tools in comparison with the evaluation on the training data. The AUC of `EpiDope` decreased by 0.065, which is within the calculated standard deviation of 0.071. The analysis of the origin of the validation sites and proteins used in this evaluation dataset shows five new families, while at the same time the distribution according to kingdom changes. This reduces the similarity with the training dataset. A lower similarity with the training data leads to somewhat limited comparability with the evaluation on the training data. At the same time, however, it indicates how well the model has generalized the epitope prediction task to the more diverse data. The AUC10% indicates the usability of all methods for practical applications. `EpiDope` outperforms all competing methods with an AUC10% of 0.089 (see Table 3). The second best method `betaturn` reaches an AUC10% of 0.069. These results indicate that `EpiDope` can predict even data that is relatively distinct from the training data and has a higher usability than competing methods.

### EpiDope output and visualization

`EpiDope` produces multiple outputs. As an easily readable and interpretable format, `EpiDope` visualizes its results in an interactive html plot using the Python `bokeh` package [21]. This allows large proteins to be displayed without impairing readability. For an example output plot see Figure 5. By default, EpiDope highlights regions of at least eight consecutive amino acids that have predicted values above the threshold of 0.818. This threshold corresponds to a 15 % recall rate with a precision of 0.635 on the training data. Furthermore the user can provide amino acid sequences (text file with one sequence per line) that are either classified as epitopes or non-epitopes to highlight them in the html plot as blue or red regions, respectively.

**Figure 5:**
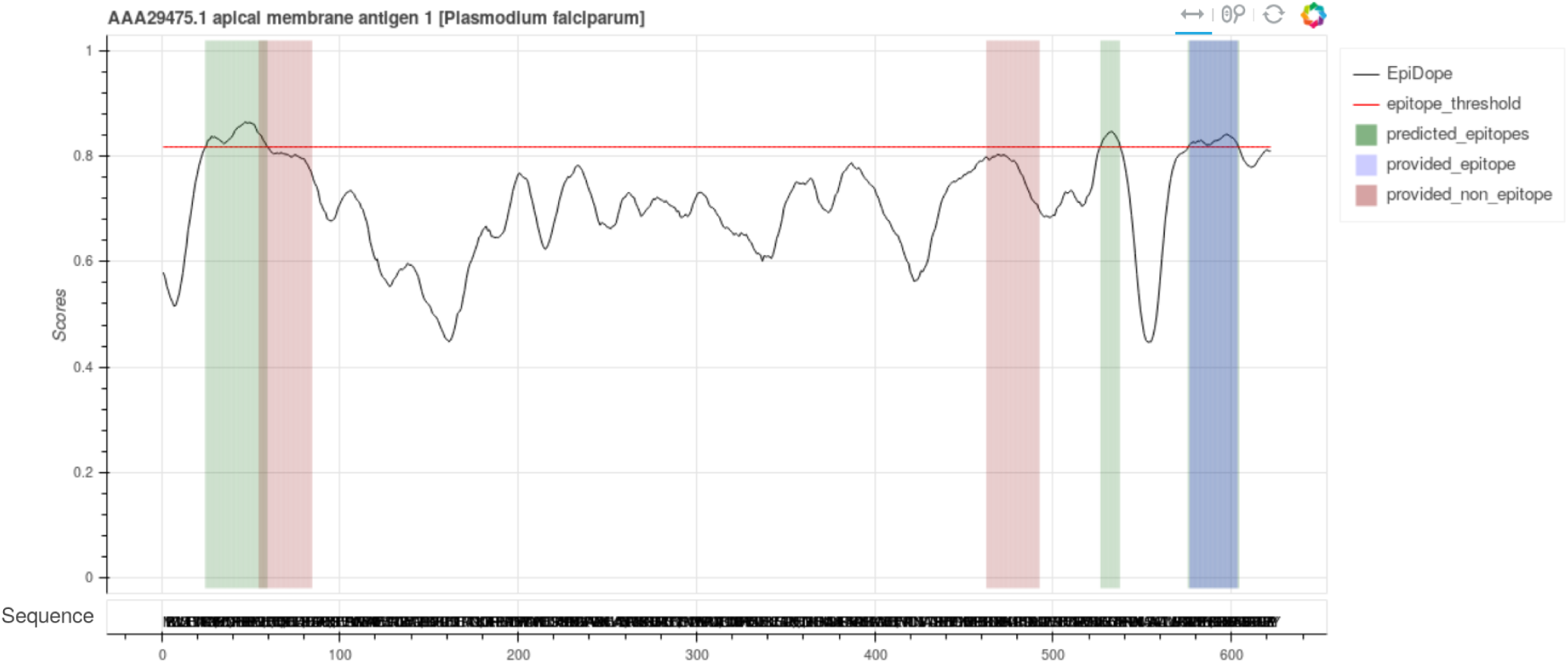
Graphical output of `EpiDope` as an interactive html plot for the apical membrane antigen 1 of *Plasmodium falciparum*. The black line displays the predicted values of `EpiDope` per position (amino acid). A higher value means that `EpiDope` certifies this region a higher chance of being an epitope. The red line is the default threshold for the predicted epitopes. The green regions are the predicted parts that are above the threshold for at least 8 consecutive amino acids. The blue and green regions are user provided regions that are known to be epitopes and non-epitopes, respectively.

Additionally, `EpiDope` produces simple computer parsable output. The file epidope_scores.csv lists the predicted score per amino acid, predicted_epitopes.csv lists all regions with a score higher than the defined threshold and predicted_epitopes_sliced.faa is a multi-fasta file of potential epitopes with a user defined size and overlap, that can be used to scan for epitopes in wet lab experiments.

## Conclusion

In this study, we have developed the linear B-cell epitope prediction tool `EpiDope` (Epitope Deep learning predictor).

While only requiring a protein’s amino acid sequence, EpiDope has been shown to be the best-performing among currently available B-cell epitope prediction tools. `EpiDope` is based on a Deep Neural Network. We trained `EpiDope` on almost 25,000 experimentally verified epitope and non-epitope regions. We have used two different datasets to compare the performance of `EpiDope` with numerous different tools, including the currently probably most used tool, `BepiPred2`. For the training data, we performed 10-fold cross-validation to ensure the reliability of the benchmarks. In addition, all proteins of a subset have a sequence identity of less than 50 % to the proteins in the other nine subsets. This ensures that all ten subsets are independent of each other. The second dataset (evaluation dataset) consists of almost 17,000 new verified epitopes and non-epitopes, which therefore have not been present in the training dataset. On both datasets `EpiDope` outperformed all competing methods. Especially for the AUC10%, which represents the performance on the practically relevant top predictions. The AUC10% was corrected by multiplying with 10 (as described in BepiPred2 [9]) and varies between 0.151 and 0.089.

The training data contains many short validated regions for which a high false-negative rate is expected [22]. Those false-negative cases result in a more conservative prediction for epitopes. However, the effect in real-world applications is minimal, as only a high true-positive and a low false-positive rate are important to reduce the lab work. These cases are best represented by AUC10%.

The high predictive power of `EpiDope` enables a much more precise search for epitopes and, thus, faster and more cost-effective development of medical treatments or diagnostic methods.

For even easier usability for a broad research community, we plan to establish an `EpiDope` Online version. Until then `EpiDope` can be downloaded from `GitHub` (github.com/mcollatz/EpiDope)or installed via `Conda`. The training datasets are available in the open science framework (osf.io/krw2j).

